# Plasma Proteomic Profiling Reveals Distinct Signatures of Chest CT Phenotypes in Sarcoidosis

**DOI:** 10.1101/2025.06.24.661339

**Authors:** Vibha Shastry, Sonia M. Leach, Brett M. Elicker, Laura L. Koth

## Abstract

**Background:** Sarcoidosis is a granulomatous disease of unknown cause with a highly variable clinical course. The inability to predict progressive inflammation, fibrosis, or both underscores the limited understanding of the underlying molecular mechanisms.

**Objective:** We aimed to identify novel protein signatures associated with distinct pulmonary phenotypes of sarcoidosis, including progressive inflammation, progressive fibrosis, and disease resolution.

**Methods:** We performed the SomaScan 11K Assay to measure more than 10,000 unique human plasma proteins and compared protein expression between chest CT-defined phenotypes using principal component analysis, differential expression, correlation analysis, and gene set enrichment analysis.

**Results:** We identified distinct proteomic signatures that differentiate progressive fibrosis from progressive nodular inflammation in sarcoidosis. Enrichment and differential expression analyses revealed that progressive fibrosis was associated with epithelial–mesenchymal transition pathways, while progressive nodular disease was linked to mTORC1 and MYC signaling, as well as metabolic activation. Additionally, expression of 44 proteins correlated moderately to strongly with thoracic lymph node enlargement, suggesting that lymph node– driven immune activity may be a major source of circulating proteomic signals.

**Conclusions:** This study leverages a unique longitudinal imaging approach to define extreme pulmonary phenotypes based on serial chest CT scoring, enabling the discovery of proteomic signals linked to distinct trajectories of sarcoidosis progression. Once validated, these findings could inform the development of blood-based biomarkers for disease stratification, monitoring, and therapeutic targeting in sarcoidosis.

## Introduction

Sarcoidosis is a multi-organ, immune-mediated granulomatous disease of unknown origin that most commonly affects the lungs and exhibits a highly variable clinical course—from spontaneous resolution to chronic inflammation, fibrosis or both. This heterogeneity complicates clinical management and trial design, as no robust, noninvasive biomarkers currently exist to distinguish these clinical trajectories. Improved patient stratification could improve clinical resource utilization, guide treatment decisions, and reduce clinical trial size. Proteomic profiling is a powerful tool for uncovering circulating biomarkers that reflect disease biology, offering a real-time readout of systemic immune activity, signaling pathways, and post-translational modifications that may be missed by genomic or transcriptomic approaches. In this study, we applied high-throughput plasma proteomics in pulmonary sarcoidosis to identify biomarkers associated with distinct CT-defined phenotypes.

We anchored biomarker discovery to longitudinal CT scans scored for fibrosis, nodular inflammation, and lymphadenopathy. Blood samples were selected from time points reflecting each subject’s most extreme CT phenotype—progressive fibrosis, progressive inflammation, or radiographic resolution. This integrative approach enabled us to link proteomic signatures to meaningful disease trajectories.

Our analysis identified distinct protein profiles associated with progressive fibrosis and inflammation. While sampling occurred concurrently with imaging, limiting predictive conclusions, these signatures provide insight into sarcoidosis immunopathology and may serve as prognostic markers pending future validation. Together, these findings underscore the value of linking proteomics with radiographic phenotypes to advance biomarker discovery in sarcoidosis.

## Methods

### Cohort Description and CT analysis

The longitudinal cohort study design and study procedures have been previously described.^1^ Participants with pulmonary sarcoidosis and healthy control subjects were recruited from the San Francisco Bay Area. Enrollment and follow-up study visits occurred at 6-to-12-month intervals between January 1, 2010, to December 31, 2021. Longitudinal chest CT scans were visually scored by an expert thoracic radiologist who was blinded to clinical data. The mean extent of reticular abnormality, honeycombing, and nodular disease was scored to the nearest 5% in three zones in each lung as previously described^2^ to produce semiquantitative scores. The severity of traction bronchiectasis was scored as previously described.^3^ In brief, severity of traction bronchiectasis within each of the five lobes was graded by comparing the diameter of the airway with that of the adjacent pulmonary artery using a 4-point scale (i.e., 0-3), and scores from the five lobes were summed to yield an overall score (range, 0 to 15): score 0, no traction bronchiectasis; score 1, traction bronchiectasis present but mild in degree (comparable diameter with the artery); score 2, moderately severe traction bronchiectasis (up to twice the diameter of the artery); and score 3, severe traction bronchiectasis (more than twice the diameter of the artery).^3^ The largest lymph node for each station (paratracheal, subcarinal and AP window) was measured using the short-axis diameter in millimeters. Size of ≤10 mm was considered normal.

### Case Control Study Design of CT Features

Longitudinal chest CT scan features were used to create three phenotype groupings based on visual scoring. These groups consisted of sarcoidosis participants with evidence of (1) progressive CT fibrosis over time defined by worsening of extent of reticulation or honeycombing or severity of traction bronchiectasis or any combination of these features; the presence of nodular inflammation was allowed; (2) progressive nodular disease over time with no or very little evidence of fibrosis; (3) remission of all or most of the CT features of disease present at baseline. Plasma samples from a healthy control group were included in the analysis but these participants did not undergo CT scan imaging. The plasma samples selected for protein analysis were from a calendar date as close as possible to the chest CT scan that was used to define the CT phenotype groupings.

### Somalogics Proteomics Assay

SomaLogic Inc. applied the SomaScan 11K Assay measuring more than 10,000 unique human proteins to previously unthawed human plasma samples that were stored at −30 °C. The SOMAmer-based assay allows for quantitative transformation of the protein epitope amount into a specific SOMAmer-based DNA signal. Fluorescent, single-stranded DNA-based SOMAmer reagents interact and bind to target molecules in the plasma, forming SOMAmer-protein complexes. Unbound proteins are removed, followed by biotinylation of SOMAmer-protein complexes. Addition of polyanionic competitors and wash steps results in specific retention of target proteins, which are then quantified using DNA-hybridization microarrays. Readouts are reported in relative fluorescent units (RFU), which are directly proportional to the amount of target protein epitope in the plasma sample. Plasma RFU data were processed using SomaLogic’s standard multi-step normalization pipeline, which applies hybridization control normalization, intraplate median normalization, plate scaling and calibration, and adaptive normalization to a reference distribution to remove both within-plate and between-run technical variation.

### Statistical Analyses

Continuous variables were summarized using means and standard deviations, and categorical variables as frequencies and percentages. Group comparisons were performed using one-way ANOVA for continuous variables and Pearson’s chi-square test for categorical variables, as implemented by the dtable command in Stata. Principal component analysis (PCA) was performed using the R stats package and the rstatix package was used to perform Kruskal-Wallis test followed by Dunn’s test on PCA data. The SomaData IO R package was used to calculate the estimated limit of detection (eLOD) per SOMAmer analyte and carry out differential expression analyses between progressive fibrosis and nodular CT phenotypes, with false discovery rate (FDR) calculated using the Benjamin-Hochberg procedure. An FDR of < 0.05 or p-value <0.05 was considered statistically significant as appropriate for the analysis. Gene Set Enrichment Analysis (GSEA) was conducted using the fgsea R package with the Hallmark gene set collection ^4^ from MSigDB. Enrichment statistics were calculated using the pre-ranked GSEA method, with normalized enrichment scores (NES) and false discovery rate (FDR) q-values provided for each pathway. Statistical and graphical analyses were carried out using R software version 4.5.0 (4-11-2025) and STATA v18 (StataCorp, College Station, TX, USA).

## Results

### Cohort / Sample Description

Table 1 summarizes the clinical characteristics of the cohort stratified by CT phenotype group. There were no statistically significant differences across the CT phenotype groups with respect to age, sex, race, tobacco history, body mass index or use of immunosuppression at the blood plasma draw date. In terms of clinical laboratory testing, there were no statistically significant differences in the levels of serum biomarkers such as C-reactive protein, sedimentation rate (ESR), or angiotensin converting enzyme (ACE), but there was a difference in the absolute lymphocyte counts with the fibrosis group having the lowest count compared to the other groups. Comparing additional clinical data within the sarcoidosis participants, there were no statistically significant differences in the total number of involved organs with sarcoidosis. The fibrosis group had the lowest mean values for lung function parameters.

**Table 1:** Clinical Characteristics at Date of Plasma Sampling*.

| Table 1: Clinical Characteristics at Date of Plasma Sampling* |  |  |  |  |  |
| --- | --- | --- | --- | --- | --- |
|  | CT Phenotype |  |  |  |  |
|  | Healthy Control | Resolved CT Features | Progressive Nodular | Progressive Fibrosis | P-Value |
| N | 5 | 5 | 15 | 15 |  |
| Age (yr) | 52 (4.4) | 47 (8.9) | 49 (12.1) | 53 (10.1) | 0.583 |
| Sex (count, %) |  |  |  |  |  |
| Male | 2 (40%) | 0 (0%) | 10 (67%) | 6 (40%) | 0.068 |
| Female | 3 (60%) | 5 (100%) | 5 (33%) | 9 (60%) |  |
| Race (count, %) |  |  |  |  |  |
| White | 3 (60%) | 3 (60%) | 12 (80%) | 10 (67%) | 0.625 |
| Black | 2 (40%) | 1 (20%) | 1 (7%) | 2 (13%) |  |
| Other | 0 (0%) | 1 (20%) | 2 (13%) | 3 (20%) |  |
| Latino (count, %) |  |  |  |  |  |
| Yes | 0 (0%) | 0 (0%) | 3 (20%) | 1 (7%) | 0.397 |
| Smoking History (count, %) |  |  |  |  |  |
| Yes | 2 (40%) | 1 (20%) | 7 (47%) | 8 (53%) | 0.626 |
| Pack-year Smoking History | 0.2 (0.3) | 3 (6.7) | 3 (6.5) | 3 (4.2) | 0.792 |
| BMI | 29 (5.0) | 30 (6.3) | 30 (6.5) | 26 (2.9) | 0.152 |
| Taking Immunosuppression on Blood Draw Date (count, %) |  |  |  |  |  |
| Yes | 0 (0%) | 0 (0%) | 3 (20%) | 5 (33%) | 0.244 |

\*all values presented as mean ± SD unless otherwise indicated

**Table 2:** Clinical Laboratory Blood Test Results at Date of Plasma Sampling*.

|  |  |  |  |  |  |
| --- | --- | --- | --- | --- | --- |
|  | CT Phenotype |  |  |  |  |
|  | Healthy Control | Resolved CT Features | Progressive Nodular | Progressive Fibrosis | P-Value |
| N | 5 | 5 | 15 | 15 |  |
| Sedimentation Rate (ESR) (mm/h) | 9.6 (6.3) | 10.3 (5.4) | 9.7 (12.4) | 13.8 (16.8) | 0.869 |
| C-Reactive Protein (mg/L) | 3.8 (2.4) | 4.1 (2.9) | 4.1 (2.5) | 11.1 (21.9) | 0.499 |
| Angiotensin Converting Enzyme (U/L) | 41 (12.5) | 28 (2.8) | 80 (69.5) | 54 (34.9) | 0.332 |
| Absolute Lymphocytes ($\times 10^9/L$) | 1.96 (0.4) | 1.73 (0.7) | 1.28 (0.5) | 1.11 (0.5) | 0.009 |

\*all values presented as mean ± SD

**Table 3:** Clinical Features at Date of Plasma Sampling*.

| Table 3: Clinical Features at Date of Plasma Sampling* |  |  |  |  |
| --- | --- | --- | --- | --- |
|  | CT Phenotype |  |  |  |
|  | Resolved CT Features | Progressive Nodular | Progressive Fibrosis | P-Value |
| N | 5 | 15 | 15 |  |
| 4 or more organs involved (count, %) |  |  |  |  |
| Yes | 3 (60%) | 6 (40%) | 4 (27%) | 0.391 |
| FEV1/FVC | 0.79 (0.06) | 0.74 (0.08) | 0.68 (0.07) | 0.026 |
| FEV1 z-score | 0.347 (0.5) | -0.395 (1.3) | -1.183 (1.0) | 0.033 |
| FEV1 %pred | 105 (7.8) | 95 (19.9) | 80 (18.1) | 0.024 |
| FVC z-score | 0.533 (0.7) | 0.168 (1.3) | -0.396 (1.0) | 0.229 |
| FVC %pred | 108 (10.1) | 103 (19.7) | 93 (16.9) | 0.191 |
| DLCO z-score | -0.416 (1.552) | -1.552 (1.276) | -1.662 (2.177) | 0.574 |
| DLCO %pred | 95 (21.6) | 80 (14.9) | 79 (25.3) | 0.526 |

\*all values presented as mean ± SD unless otherwise indicated

Table 4 presents the summary data for the chest CT scan features that were present at the time of plasma collection. The resolved CT feature group had no evidence of reticulation, traction bronchiectasis and had the lowest mean average size of mediastinal lymph nodes. The progressive nodular group was characterized by the highest mean percentage of lung involved with nodular opacities, and the highest mean mediastinal lymph node size. Among participants in the progressive nodular group, only one exhibited lung parenchymal fibrosis, with 10% reticulation and the highest observed traction bronchiectasis score of “3”. Additionally, two other participants in this group had evidence of traction bronchiectasis without associated parenchymal fibrosis. The progressive fibrosis group had the highest average level of reticulation and traction bronchiectasis. There was no difference in the average number of years between the baseline and the last follow-up chest CT.

**Table 4:** Chest CT Scan Features at Date of Plasma Sampling*.

| Table 4: Chest CT Scan Features at Date of Plasma Sampling* |  |  |  |  |
| --- | --- | --- | --- | --- |
|  | CT Phenotype |  |  |  |
|  | Resolved CT Features | Progressive Nodular | Progressive Fibrosis | P-Value |
| N | 5 | 15 | 15 |  |
| Percent of Lung Involved with Nodules | 4 (6.5) | 41 (19.0) | 23 (23.8) | 0.003 |
| Percent of Lung Involved with Reticulation | 0 (0.0) | 1 (2.6) | 18 (16.4) | <0.001 |
| Traction Bronchiectasis Score (count, %) |  |  |  |  |
| 0 | 5 (100%) | 12 (80%) | 1 (7%) | 0.006 |
| 2 | 0 (0%) | 2 (13%) | 4 (27%) |  |
| 3 | 0 (0%) | 1 (7%) | 1 (7%) |  |
| 4 | 0 (0%) | 0 (0%) | 5 (33%) |  |
| 8 | 0 (0%) | 0 (0%) | 1 (7%) |  |
| 9 | 0 (0%) | 0 (0%) | 3 (20%) |  |
| Lymph Node Size AP Window (mm) | 5 (2.1) | 11 (4.0) | 7 (2.6) | <0.001 |
| Lymph Node Size Paratracheal (mm) | 7 (4.2) | 13 (5.4) | 9 (4.7) | 0.048 |
| Lymph Node Size Subcarinal (mm) | 8 (4.3) | 15 (4.0) | 11 (4.9) | 0.007 |
| Years between baseline and last follow-up CT scan | 4.6 (2.6) | 6.3 (4.2) | 5.7 (3.9) | 0.686 |

\*all values presented as mean ± SD unless otherwise indicated

### Global Data Overview / QC/ PCA

We used the SomaScan assay to measure 10,760 plasma proteins across 35 patients with sarcoidosis and 5 healthy unaffected volunteers. We performed principal component analysis (PCA) and identified four sample outliers that were dropped in subsequent analyses. Three out of four of these samples were also flagged by the SomaLogics quality control processing steps. Of the four outliers, three of them belonged to the nodular group, and one belonged to the resolving group. To ensure confidence in selecting high abundance proteins, a filtering parameter was created representing the signal to noise (STN) ratio, obtained by dividing the median RFU per analyte by the estimated limit of detection (eLOD) of the assay buffer controls as recommended by the assay manufacturer. A plot depicting the distribution of STN ratios indicated a clear inflection point around a STN ratio of 31.8, where the signal to noise ratio increased dramatically (**Supplemental Figure 1**). For our downstream analyses, we used varying numbers of proteins that were observed as having a STN ratio greater than this inflection point. We confirmed that this filtering strategy retained analytes with the highest variance across the samples (**Supplemental Figure 2**).

Figure 1A depicts the PCA analysis after dropping the four outliers and using the previously mentioned STN filtering parameter. This result suggested reasonable separation of progressive fibrosis and nodular CT phenotypes, although some overlap between the two disease clusters remained. Principal component (PC) 1 explained 27% of the variance in the scaled and centered RFU values, while principal component 2 explained 12%. A plot comparing PC1 scores across all phenotypes showed a statistically significant difference between progressive fibrosis and nodular groups (Figures 1B). No statistically significant differences in PC1 or PC2 scores between other disease phenotypes were observed (Figures 1B-C). These results suggested differential regulation between progressive fibrosis and nodular CT phenotypes and motivated additional analyses.

**Figure 1:**
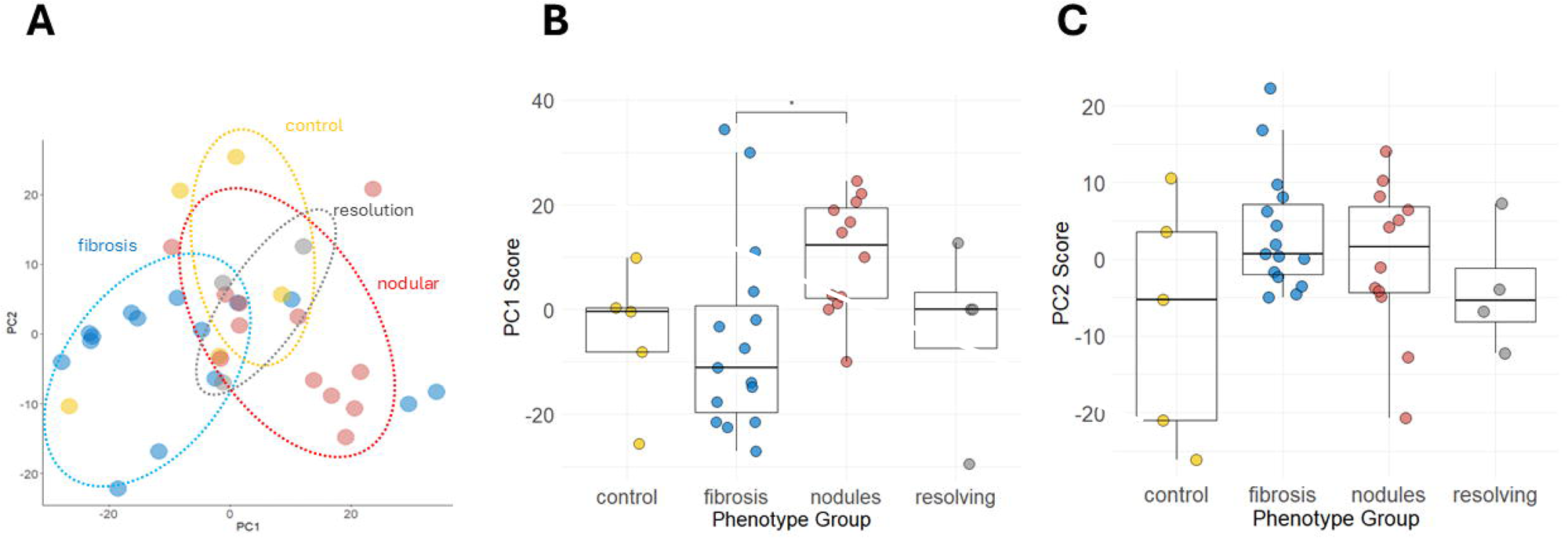
Principal Component Analysis (PCA) of proteomics data generated from plasma samples. (A) PCA results are displayed after removal of four sample outliers. The proteomics data included in this analysis consisted of analytes with a signal to noise ratio of greater or equal to 31.8 and consisted of 2,006 analytes. Plots in panel B and C show the individual principal component (PC) scores for PC1 and PC2 plotted against the phenotype group. A Kruskal-Wallis rank sum test and post-estimation Dunn’s test with Bonferroni correction was performed to evaluate median PC score differences across the groups. *p-value of <0.05

### Exploratory Analysis via Unsupervised Clustering

To further visualize the difference in the range of protein expression across groups, a global comparison of protein expression was performed using unsupervised hierarchal clustering analysis. This approach used the Ward D2 minimum variance clustering method with correlation distance and included the top 2,006 analytes as determined by STN ratio (analytes above the inflection point from Supplemental Figure 1). Figure 2A displays the clustering analysis with the inclusion of all three CT phenotypes and healthy controls. One cluster contained mostly participants with the progressive fibrosis phenotype (13/20), while another cluster largely contained those with progressive nodular phenotype (8/16). When comparing proteins, three distinct clusters were observed, each containing 554, 869, and 583 analytes respectively. The above observation prompted performing another iteration of cluster analysis, this time with a focus on only the progressive fibrosis and nodular phenotypes. Figure 2B displays this result showing two distinct protein clusters, with 1169 and 869 proteins contained in each cluster.

**Figure 2:**
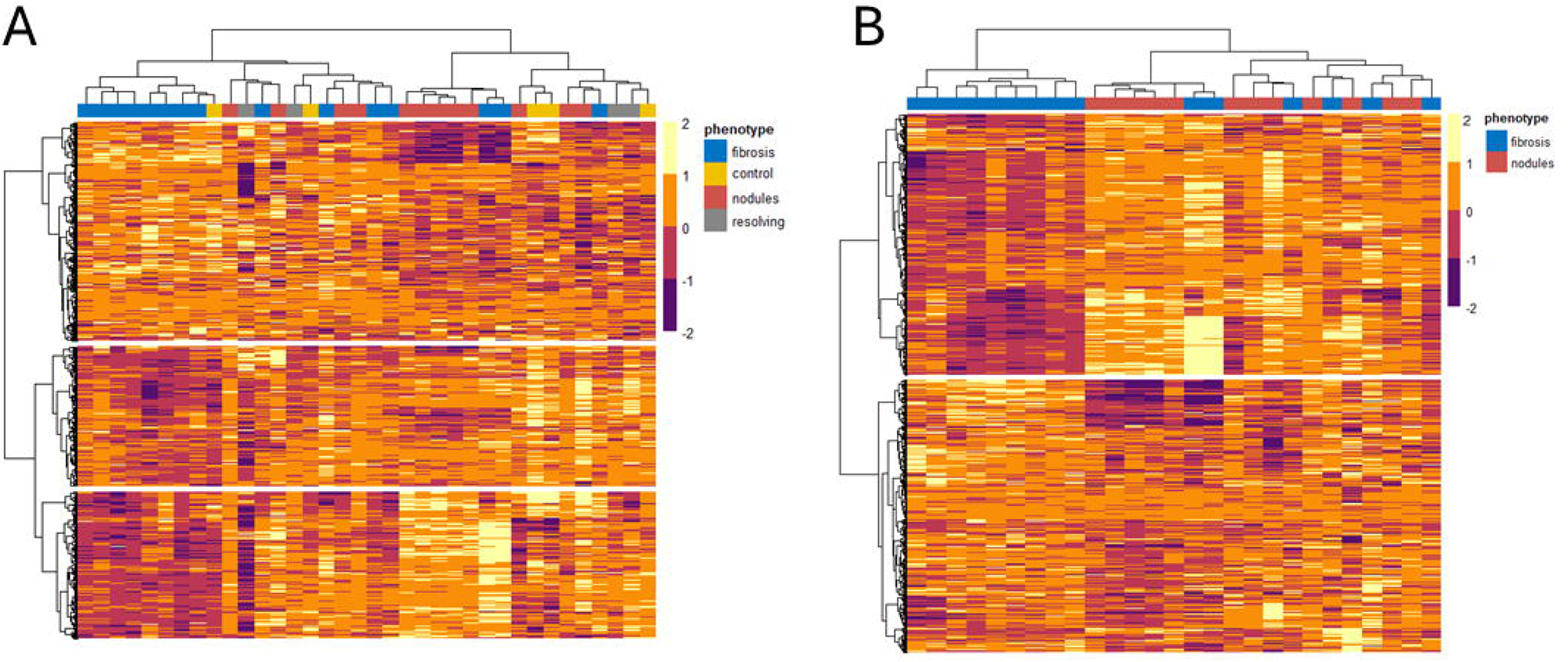
Heatmap of protein relative expression across samples. Unsupervised hierarchical clustering was applied using Ward’s D2 linkage method and correlation distance as the dissimilarity metric. Rows represent proteins; columns represent individual samples, grouped by cluster membership. Input for analysis included the top 2,006 analytes as determined by signal to noise ratio > 31.8 (analytes above the inflection point from Supplemental Figure 1). Figure 2A displays the clustering analysis with inclusion of all three CT phenotypes and healthy controls. When comparing proteins, three distinct clusters were observed, each containing 554, 869, and 583 analytes, respectively. Figure 2B repeated the same analysis but only included the progressive fibrosis and nodular phenotypes. This result yielded two distinct protein clusters, with 1169 and 869 proteins contained in each cluster.

### Differential Expression Results

Given that the progressive fibrosis and nodular CT phenotypes exhibited potential differences in protein expression based on PCA results and unsupervised hierarchal clustering, a comparison between both CT phenotypes was visualized using a volcano plot (Figure 3). The volcano plot depicts differentially expressed proteins when comparing progressive fibrosis to nodular phenotype groups. The analytes with marker colors represent those with both an FDR cutoff less than 10% and a log 2-fold change (log2FC) greater than 0.6 or less than −0.6 (i.e., an absolute FC of 1.5). The analytes above the horizontal line were those with an FDR < 0.05. A greater proportion of down- to up-regulated proteins was observed in the progressive fibrosis phenotype compared to the nodular phenotype. The significantly downregulated proteins in the fibrosis phenotype, ANXA2, MVP, and GBP1, had a log2FC <-1.5 and FDR < 0.05. Another protein, CMPK1, while having FDR < 0.05, had a log2FC >-1.5. Sixty-five analytes had FDR = 0.06 (**Supplemental Table 1**), and when raising the false discovery rate threshold to 0.1 or 10%, the number of differentially expressed proteins increased to 186. Other analytes that were upregulated in the fibrotic CT phenotype compared with the nodular group with an FDR ≤ 0.1 were notable given their proposed roles in fibrotic pathophysiology. These analytes include CYP24A1, FBXO22, BCL9, Matrillin-2, and Fibulin-7. These findings are notable given the modest sample size and highlight the potential of quantitative CT-derived phenotyping as a sensitive approach for uncovering underlying biological mechanisms—an approach that remains underutilized in sarcoidosis research. No differentially expressed proteins were detected when comparing other disease phenotypes, which could be attributed to the relatively smaller number of patients assigned those phenotype groupings.

**Figure 3:**
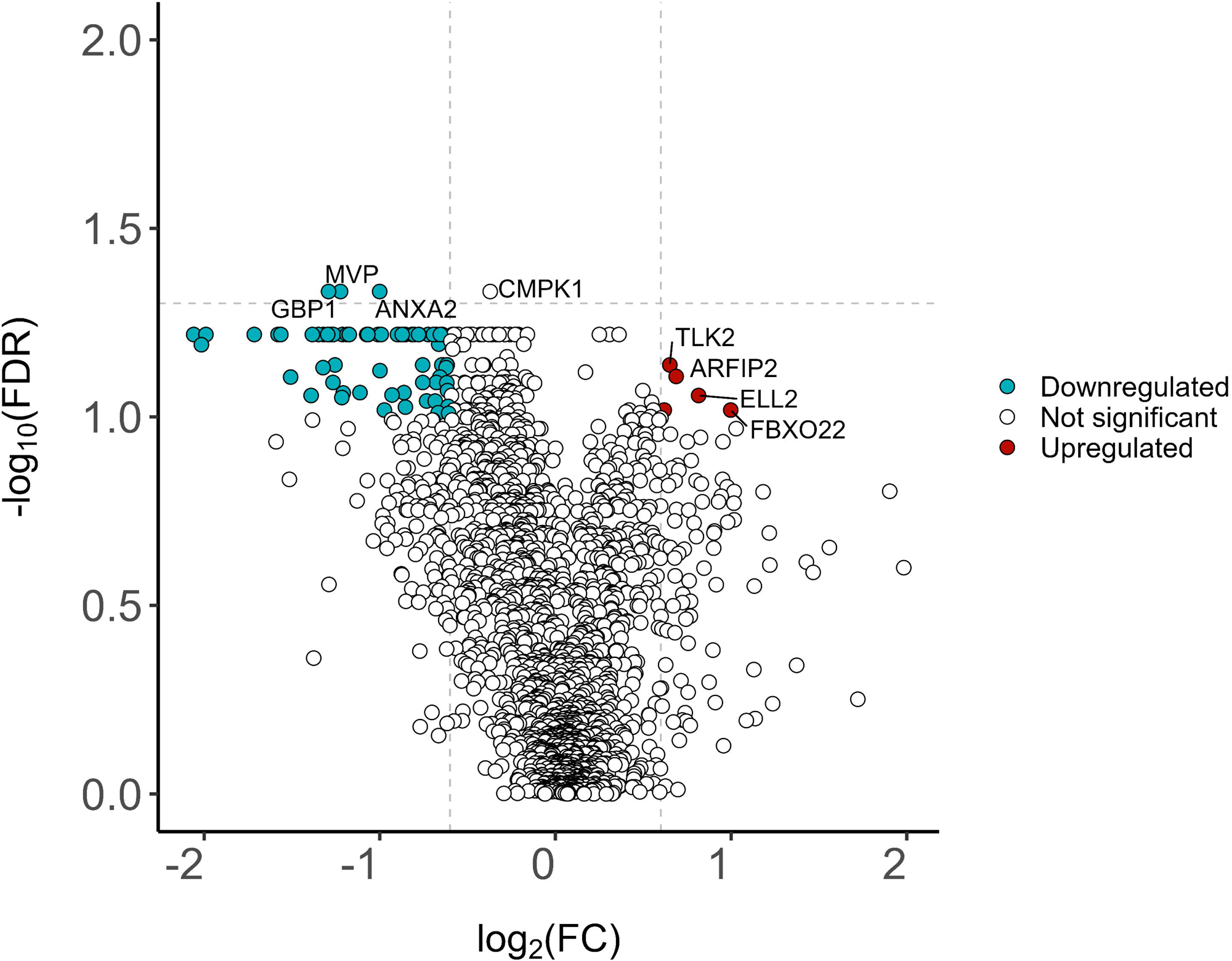
Volcano plot comparing the protein expression in the progressive fibrosis to nodular CT phenotypes. The top 2,006 analytes by signal to noise ratio were evaluated. Proteins were classified as upregulated (colored red) or downregulated (colored blue) in progressive fibrosis based on having a log2fold change (log2FC) threshold of +/− 0.6 (equivalent to a fold change +/− 1.5 and indicated by the vertical dashed lines) and –log10 false discovery rate (-log10FDR) = 1 (equivalent to FDR=0.1 and indicated by the horizontal dashed line). Four downregulated proteins (GBP, MVP, ANXA2, CMPK1) met a stricter FDR significance threshold of 0.05, though CMPK failed to meet the 1.5-fold-change threshold for significance.

### Enrichment Analyses

To examine coordinated differences of protein expression between the progressive fibrosis and nodular phenotypes, we performed gene set enrichment analysis (GSEA). The entire set of analytes were mapped to gene identifiers and subsequently ranked by p-value multiplied by the sign of log2 FC. The gene sets that were enriched with an FDR <0.05 in the nodular phenotype group were pathways associated with mTORC1 signaling, oxidative phosphorylation, adipogenesis, fatty acid metabolism, and MYC signaling which are illustrated in Figures 4A**-**4D. The epithelial mesenchymal transition (EMT) gene set with an FDR < 0.05 was enriched in the progressive fibrosis phenotype. (Figure 4E).

**Figure 4:**
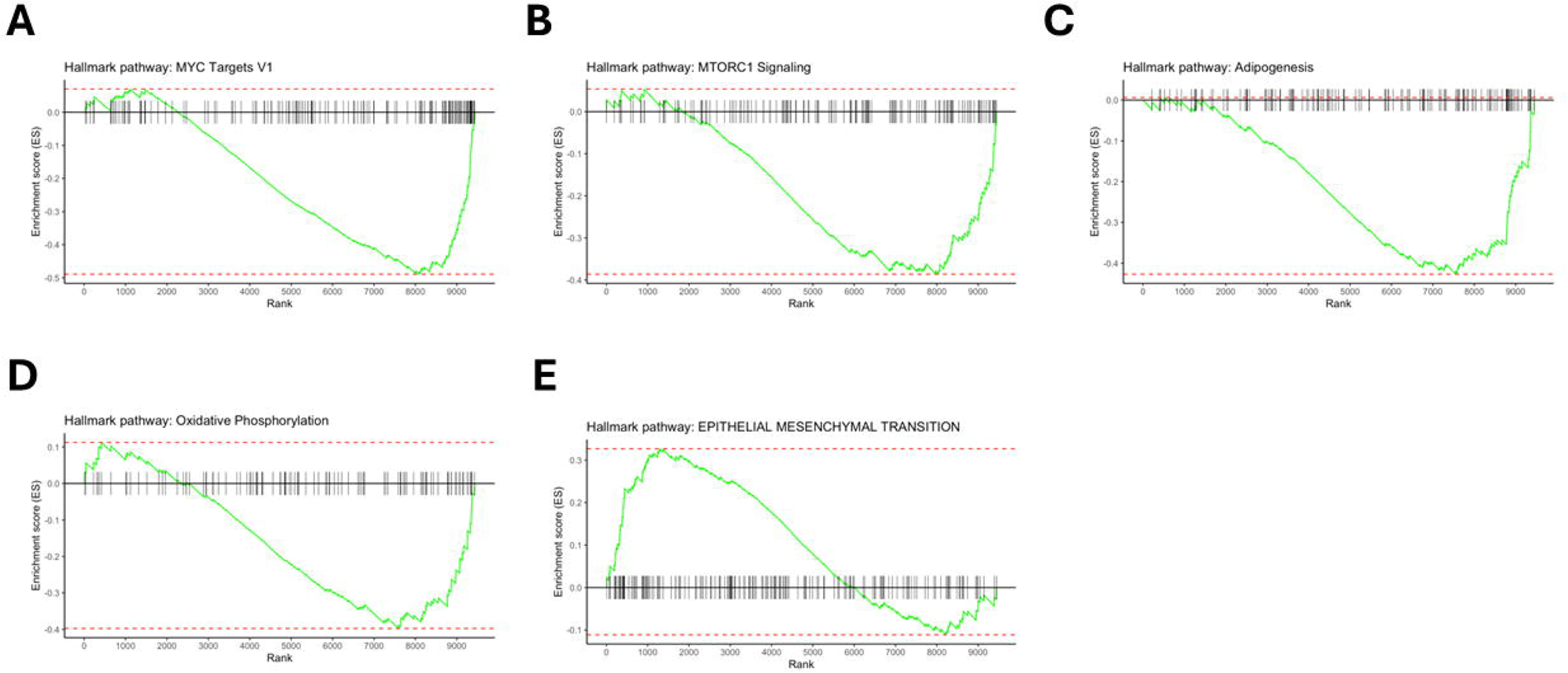
Gene set enrichment analysis (GSEA) performed on proteins ranked by differential expression between progressive fibrosis and nodular CT phenotypes. Enrichment results using the Hallmark gene set collection for the (A) MYC Targets V1 gene set, and (B) MTORC1 Signaling pathway, (C) Adipogenesis, (D) Oxidative Phosphorylation, and (E) Epithelial Mesenchymal Transition are displayed. Gene sets shown in A-D were enriched in the progressive nodular CT phenotype group compared to progressive fibrosis while the gene set for E was enriched in the progressive fibrosis compared to the nodular group.

### Correlation of Protein Expression with Chest CT Features and Blood Biomarkers

To understand how the top 70 differentially expressed proteins between progressive fibrosis and nodular CT phenotypes compared to specific chest CT features, we performed unsupervised hierarchal clustering (Figure 5). The plot illustrates the striking difference in expression of these protein analytes when comparing lung reticulation vs nodular opacities. Even more striking and unexpected was the moderate to high correlations between the size of the lymph nodes and the protein expression as observed in a cluster of 44 analytes. One implication of this finding is that the profile of circulating plasma proteins may be more influenced by the immune activity occurring within lymph nodes than from within lung parenchymal nodules.

**Figure 5:**
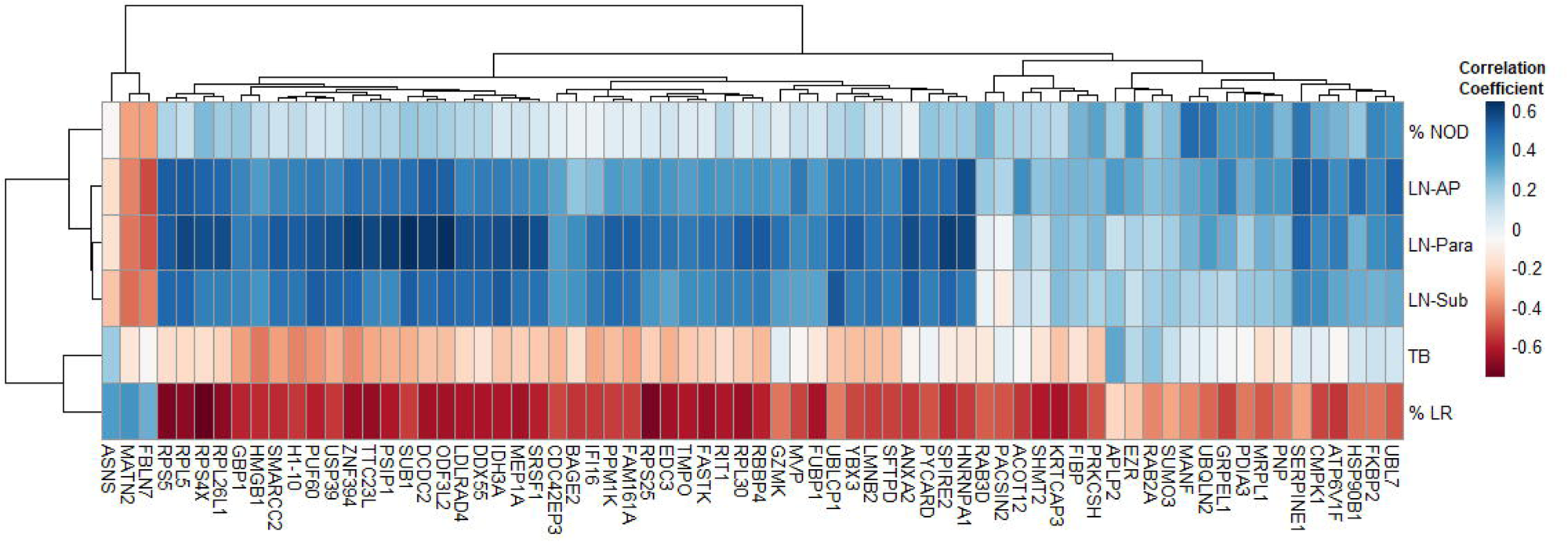
Heatmap with hierarchical clustering of differentially expressed analytes between progressive fibrosis and nodular CT phenotypes. Analytes were clustered based on correlation coefficients calculated between analyte relative fluorescence units and selected chest CT features, which were visually scored by an expert radiologist. CT features, listed top-to-bottom, include the percentage of nodular opacities (%NOD), followed by the size of the largest lymph node in three mediastinal regions—aortopulmonary window (LN-AP), paratracheal (LN-Para), and subcarinal (LN-Sub), severity of traction bronchiectasis (TB) and percentage of lung reticulation (%LR). The FDR threshold used to select these analytes was FDR ≤ 0.06.

The RFU values of differentially expressed analytes were also correlated with clinically available biomarker levels measured at the same study visit as the plasma sample used in the proteomic analysis. Figure 6 again displays the results of unsupervised hierarchal clustering analysis and reveals weak correlations between serum levels of CD25 (aka soluble IL-2Ra), angiotensin converting enzyme (ACE), and lymphocyte count using the same list of differentially expressed analytes between progressive fibrosis vs nodular phenotypes. In contrast, C-Reactive Protein (CRP) and lysozyme showed weak to moderate negative correlations with most of these analytes. A small group of analytes were positively correlated with CRP of which two, UBL7^5^ and UBQLN2^6^, have been linked to IL-6 related inflammatory pathways.

**Figure 6:**
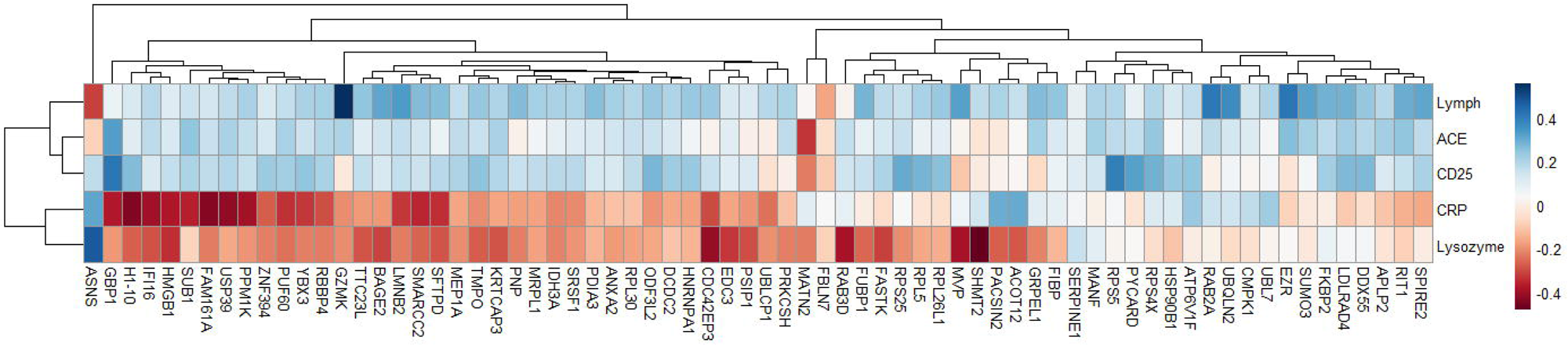
Heatmap with hierarchical clustering of differentially expressed analytes between progressive fibrosis and nodular CT phenotypes. Analytes were clustered based on correlation coefficients calculated between analyte relative fluorescence units and blood biomarker levels that were measured at the same time point used for proteomics measurements. The blood tests presented from top to bottom include absolute lymphocytes (10^9^/L) (Lymph), Angiotensin Converting Enzyme (U/L) (ACE), soluble IL2RA (pg/mL) (CD25), C-Reactive Protein (mg/L) (CRP), and lysozyme (ng/mL). The FDR threshold used for these analytes was FDR ≤ 0.06.

## Discussion

This study highlights the power of integrating high-resolution pulmonary phenotyping from longitudinal CT imaging with deep plasma proteomics. By leveraging a small but rigorously characterized cohort, we identified biologically plausible protein signatures linked to distinct CT-defined sarcoidosis phenotypes. Specifically, we found enrichment of protein signatures related to metabolic activity in the progressive nodular group, while enrichment of proteins related to epithelial-mesenchymal transition and other fibrotic mechanisms were found in the progressive fibrosis group.

From our differential protein analysis, we identified three proteins that were significantly downregulated in the progressive fibrosis group that play plausible mechanistic roles in the development of lung fibrosis. These proteins include ANXA2, a member of the annexin-family of calcium-dependent cytosolic proteins involved in phospholipid binding and plasma membrane repair. Dysregulation of ANXA2 has been previously associated with impaired fibrinolysis ^7^ and may promote fibrin accumulation, leading to increased collagen and scar formation. Support for ANXA2’s role in sarcoidosis-related fibrosis is strengthened by similarly decreased levels in idiopathic pulmonary fibrosis (IPF).^8^ Another highly downregulated protein in the fibrosis group was GBP1, Guanylate-binding protein 1. GBP1 plays a protective role against inflammation-induced cellular damage and dysfunction. It promotes mitophagy, which is a process that clears damaged mitochondria, and helps maintain mitochondrial function^9^. Thus, severe downregulation of GBP1 as found in the fibrotic group may be associated with a cascade of cellular events that create an environment promoting fibrosis by contributing to mitochondrial dysfunction, oxidative stress, inflammation, and cellular senescence.^9^ Other proteins like lung resistance protein (LRP), also known as major vault protein (MVP), is the main component of vaults, which are large ribonucleoprotein particles. While the exact role of vaults and MVP is not fully understood, a severe downregulation could contribute to fibrosis through its role in regulating several cellular pathways including impaired cell survival and increased apoptosis. Observed downregulation of MVP has also been previously found in lung tissues from IPF.^10^ Thus, two of the most highly downregulated proteins in the progressive fibrotic group are also downregulated in prototypical progressive fibrotic lung disease, IPF.

We also found a group of protein analytes upregulated in the progressive fibrosis group. Many of them have known or putative roles in fibrotic pathophysiology, underscoring their potential relevance in sarcoidosis-related tissue remodeling. For example, Cytochrome P450 24A1 (CYP24A1) promotes the breakdown of active vitamin D (1,25-dihydroxyvitamin D), thereby reducing vitamin D signaling, which is known to exert anti-fibrotic effects.^11–13^ Elevated expression of CYP24A1 may therefore contribute to fibrosis by limiting this protective signaling pathway. P2X purinoceptor 4 (P2RX4) is an ATP-gated ion channel that contributes to mechanosensation and inflammation, with growing evidence supporting its role in promoting fibrotic remodeling, particularly in cardiac and pulmonary contexts.^14,15^ Matrilin-2, a structural extracellular matrix (ECM) protein, facilitates cell-matrix interactions and has been associated with fibrotic tissue architecture and fibroblast migration.^16,17^ Lastly, Fibulin-7, a newer member of the fibulin ECM glycoprotein family, binds to the epidermal growth factor receptor (EGFR) and activates downstream signaling pathways, promoting fibroblast-to-myofibroblast trans-differentiation and collagen deposition in cardiac fibrosis.^17^ The convergence of these analytes in established fibrosis pathways suggests a shared biological axis that may underlie progressive fibrosis in sarcoidosis and warrants validation and further mechanistic investigation.

The pathway analysis offers more insight into the types of molecular drivers underlying the progressive nodular phenotype. The positively enriched pathways—mTORC1 signaling, oxidative phosphorylation, adipogenesis, fatty acid metabolism, and MYC signaling collectively reflect a coordinated shift toward enhanced cellular metabolism and cellular growth regulation, suggesting that metabolic reprogramming may play a central role in the pathobiology of progressive inflammatory sarcoidosis. The mTORC1 observation aligns with experimental findings showing that chronic mTORC1 activation in macrophages drives granuloma formation and proinflammatory remodeling, providing a mechanistic link between our proteomic signature and persistent granulomatous inflammation.^18^ Furthermore, the MYC transcription factor, involved in cell proliferation and growth, has also been associated with greater formation of multinuclear macrophage subsets that comprise granulomas.^19^

Pathway analyses results in the fibrosis group point to tissue remodeling pathways, such as the epithelial-mesenchymal transition (EMT), a key cellular program driving fibroblast accumulation and extracellular matrix production. Relevant EMT proteins that were differentially expressed include F-box only protein 22 (FBXO22), a component of the SCF E3 ubiquitin ligase complex, that has been shown to promote epithelial-mesenchymal transition (EMT).^20,21^ B-cell CLL/lymphoma 9 (BCL9) functions as a transcriptional co-activator in the Wnt/β-catenin pathway, which is widely implicated in fibrotic diseases through its regulation of mesenchymal cell activation and collagen deposition.^22,23^ Similarly, receptor tyrosine kinase-like orphan receptor 1 (ROR1) participates in non-canonical Wnt signaling and has been linked to profibrotic EMT in multiple organ systems, including the lung.^24,25^

Strengths of the analysis include the use of proteomic analysis which offers key advantages by capturing real-time biological activity at the protein level, including post-translational modifications not detected by transcriptomics. It enables discovery of clinically relevant biomarkers and dysregulated pathways to bridge molecular mechanisms with disease manifestations, enhancing translational insight. While these results shed greater light into biological pathways understudied in sarcoidosis, we acknowledge several important limitations including the sample size of our cohort which limits generalizability and the unknown effects of immunosuppressants on protein expression. We attempted to limit the effect of this latter issue by limiting the number of participants taking immunosuppression.

In summary, these findings highlight the potential of quantitative CT-derived phenotyping as a sensitive approach for uncovering underlying biological mechanisms—an approach that remains underutilized in sarcoidosis research. Collectively, these peripheral blood protein profile signatures found in the fibrosis and nodular groups appear to distinguish the groups and may reflect the types of underlying mechanisms occurring in tissues. As no biomarkers are currently available for progressive pulmonary fibrosis in sarcoidosis, the proteins highlighted in this study offer exciting candidates for validation studies.

## Author Contributions

L.L.K. curated the data and conceived of the study. V.S. performed data analysis, statistical modeling and figure generation. V.S., S.M.L., and L.L.K. performed the analyses interpretations. B.M.E. performed the scoring of imaging findings. All authors contributed to writing the manuscript, provided critical revisions, and approved the final version.

