## Supplemental Figures 1-2 for "Plasma Proteomic Profiling Reveals Distinct Signatures of Chest CT Phenotypes in Sarcoidosis"

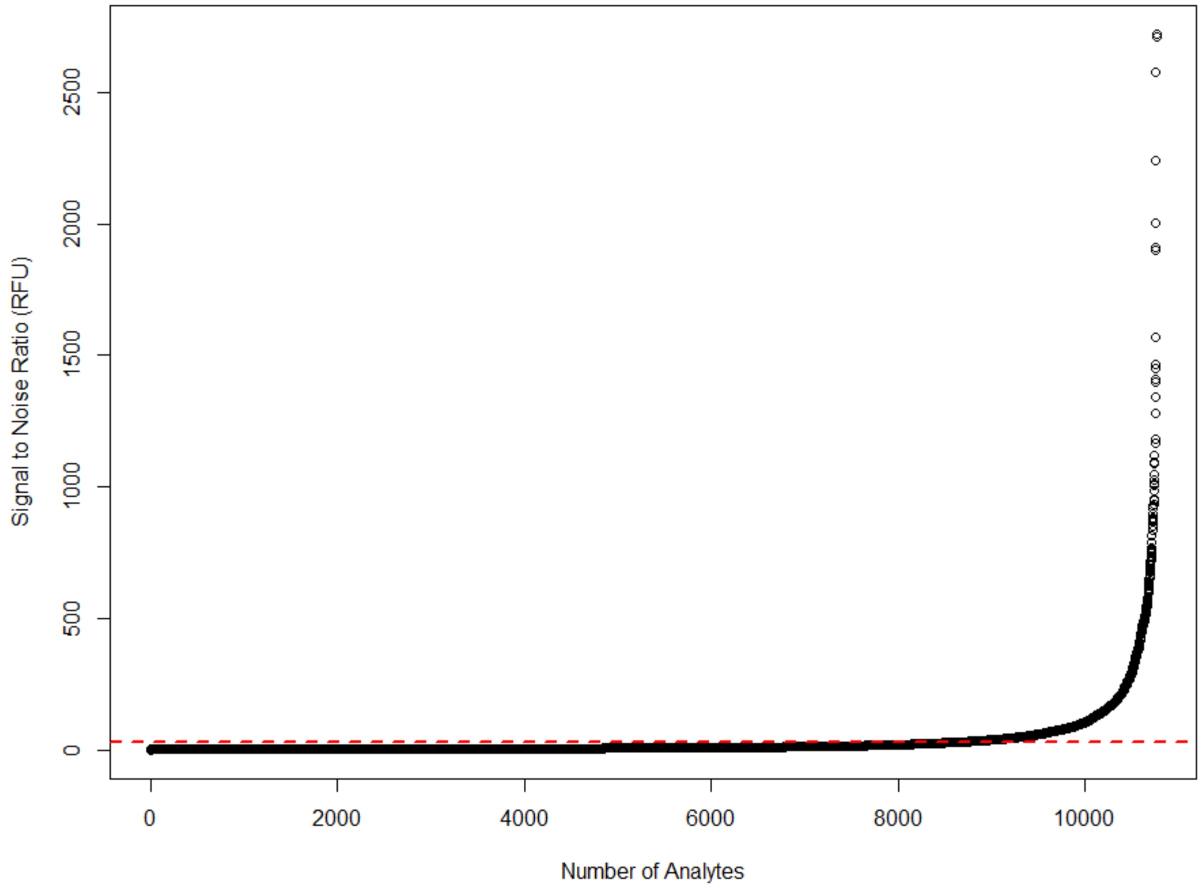


**Supplemental Figure 1:** Plot of the signal to noise parameter vs relative fluorescence units of the protein analytes. The curve shows an inflection point around a signal to noise ratio of 31.8, where the curve increases dramatically, and results in 2006 analytes with a signal to noise ratio > 31.8.

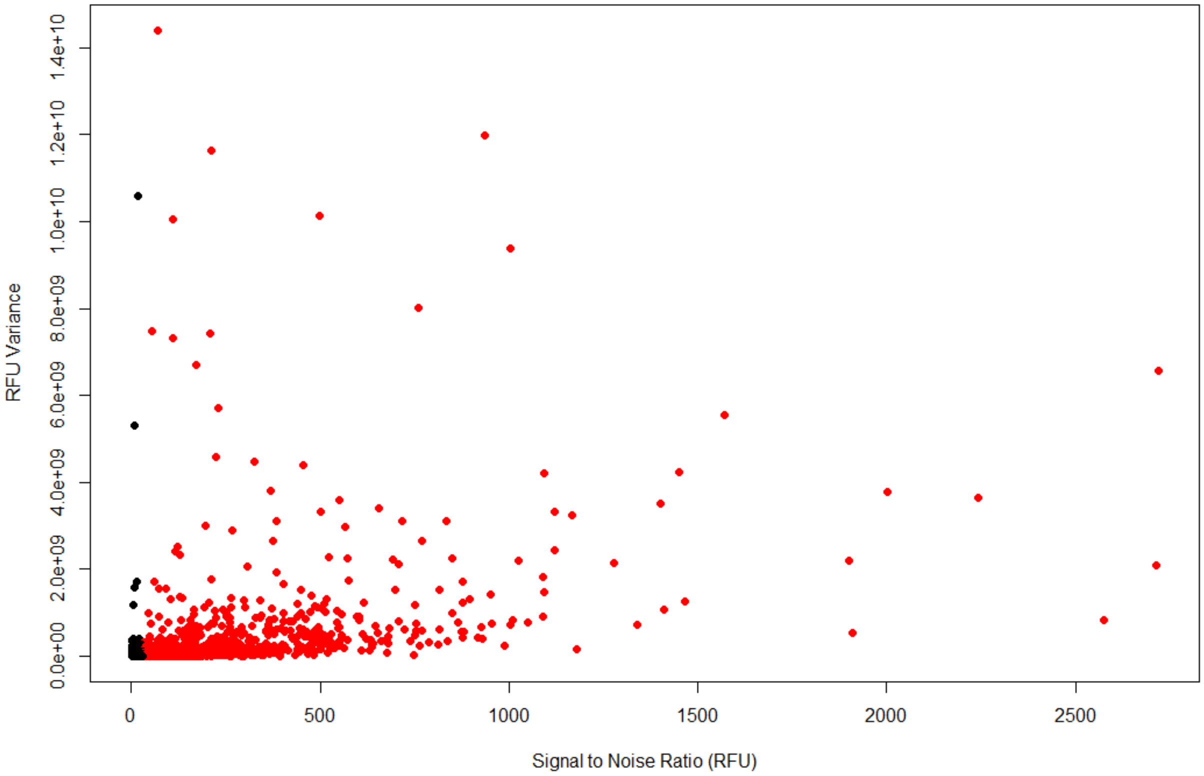


**Supplemental Figure 2**: Plot of the calculated variance versus signal to noise ratio. The red color marker indicates the 2,006 analytes with signal to noise ratio values above 31.8, the inflection point depicted in Supplemental Figure 1.

| Aptamer Name | Target | Entrez Gene Symbol | FDR | log2FC |
|---|---|---|---|---|
| seq.29660.153 | Isocitrate dehydrogenase [NAD] subunit alpha, mitochondrial | IDH3A | 0.0604733 | -1.212492 |
| seq.10575.31 | Poly(U)-binding-splicing factor PUF60 | PUF60 | 0.0604733 | -0.579354 |
| seq.19195.85 | 40S ribosomal protein S5 | RPS5 | 0.0604733 | -0.64865 |
| seq.3325.2 | Matrilin-2 | MATN2 | 0.0604733 | 0.3078188 |
| seq.31440.109 | Doublecortin domain-containing protein 2 | DCDC2 | 0.0604733 | -0.5859 |
| seq.33622.23 | Zinc finger protein 394 | ZNF394 | 0.0604733 | -0.854455 |
| seq.20217.26 | Lamin-B2 | LMNB2 | 0.0604733 | -1.277159 |
| seq.30977.31 | Enhancer of mRNA-decapping protein 3 | EDC3 | 0.0604733 | -0.724611 |
| seq.30274.3 | U4/U6.U5 tri-snRNP-associated protein 2 | USP39 | 0.0604733 | -1.579421 |
| seq.19124.9 | Ubiquitin-like domain-containing CTD phosphatase 1 | UBLCP1 | 0.0604733 | -0.477062 |
| seq.20999.12 | 60S ribosomal protein L5 | RPL5 | 0.0604733 | -1.010006 |
| seq.19808.26 | Ras-related protein Rab-3D | RAB3D | 0.0604733 | -0.816341 |
| seq.9758.17 | 40S ribosomal protein S4, X isoform | RPS4X | 0.0604733 | -0.777589 |
| seq.30501.60 | GTP-binding protein Rit1 | RIT1 | 0.0604733 | -0.892728 |
| seq.7113.1 | GrpE protein homolog 1, mitochondrial | GRPEL1 | 0.0604733 | -0.491573 |
| seq.33170.45 | Tetratricopeptide repeat protein 23-like | TTC23L | 0.0604733 | -0.680107 |
| seq.20960.47 | Far upstream element-binding protein 1 | FUBP1 | 0.0604733 | -0.499944 |
| seq.13940.19 | Gamma-interferon-inducible protein 16:Isoform 2, Hematopoietic expression, interferon-inducible nature, and nuclear localization 1 | IFI16 | 0.0604733 | -1.320821 |
| seq.23694.3 | Asparagine synthetase | ASNS | 0.0604733 | 0.3618345 |
| seq.21387.64 | Meprin A subunit alpha | MEP1A | 0.0604733 | -1.074175 |
| seq.8265.225 | Lamina-associated polypeptide 2, isoforms beta/gamma | TMPO | 0.0604733 | -0.899581 |
| seq.28164.82 | Outer dense fiber protein 3-like protein 2 | ODF3L2 | 0.0604733 | -1.065512 |
| seq.2925.9 | Plasminogen activator inhibitor 1 | SERPINE1 | 0.0604733 | -0.210774 |
| seq.16857.2 | Ras-related protein Rab-2A | RAB2A | 0.0604733 | -0.557741 |
| seq.31035.50 | Protein spire homolog 2 | SPIRE2 | 0.0604733 | -0.804764 |
| seq.29250.39 | BRG1-associated factor 170 | SMARCC2 | 0.0604733 | -1.264628 |
| seq.31015.10 | Protein phosphatase 1K, mitochondrial | PPM1K | 0.0604733 | -2.059167 |
| seq.31932.34 | Keratinocyte-associated protein 3 | KRTCAP3 | 0.0604733 | -0.368416 |
| seq.12478.15 | 60S ribosomal protein L30 | RPL30 | 0.0604733 | -0.779151 |

| | | | | |
|---|---|---|---|---|
| seq.29504.25 | Mesencephalic astrocyte-derived neurotrophic factor | MANF | 0.0604733 | -0.258117 |
| seq.9339.204 | Peptidyl-prolyl cis-trans isomerase FKBP2 | FKBP2 | 0.0604733 | -0.287633 |
| seq.24671.15 | Apoptosis-associated speck-like protein containing a CARD | PYCARD | 0.0604733 | -0.22847 |
| seq.7832.181 | Protein disulfide-isomerase A3 | PDIA3 | 0.0604733 | -0.417827 |
| seq.4414.69 | Pulmonary surfactant-associated protein D | SFTPD | 0.0604733 | -1.563344 |
| seq.20433.19 | 39S ribosomal protein L1, mitochondrial | MRPL1 | 0.0604733 | -0.677619 |
| seq.10627.87 | Amyloid-like protein 2 | APLP2 | 0.0604733 | -0.428488 |
| seq.19121.3 | Ubiquilin-2 | UBQLN2 | 0.0604733 | -0.219644 |
| seq.20137.49 | Heterogeneous nuclear ribonucleoprotein A1 | HNRNPA1 | 0.0604733 | -1.186569 |
| seq.31412.22 | Fas-activated serine/threonine kinase | FASTK | 0.0604733 | -1.175989 |
| seq.6393.63 | Endoplasmin | HSP90B1 | 0.0604733 | -0.471026 |
| seq.25048.30 | Protein kinase C and casein kinase substrate in neurons protein 2 | PACSIN2 | 0.0604733 | -0.413275 |
| seq.21583.14 | 60S ribosomal protein L26-like 1 | RPL26L1 | 0.0604733 | -0.694338 |
| seq.19236.24 | Activated RNA polymerase II transcriptional coactivator p15 | SUB1 | 0.0604733 | -1.348913 |
| seq.25219.17 | V-type proton ATPase subunit F | ATP6V1F | 0.0604733 | -0.50372 |
| seq.9753.17 | Ezrin | EZR | 0.0604733 | -0.229241 |
| seq.33227.149 | Protein FAM161A | FAM161A | 0.0604733 | -1.714632 |
| seq.15435.4 | Purine nucleoside phosphorylase | PNP | 0.0604733 | -0.55272 |
| seq.9545.156 | Granzyme K | GZMK | 0.0604733 | -1.173976 |
| seq.26209.2 | Splicing factor, arginine/serine-rich 1 | SRSF1 | 0.0604733 | -0.99152 |
| seq.19169.88 | Acidic fibroblast growth factor intracellular-binding protein | FIBP | 0.0604733 | -0.370558 |
| seq.23243.120 | 40S ribosomal protein S25 | RPS25 | 0.0604733 | -0.33538 |
| seq.17176.13 | PC4 and SFRS1-interacting protein | PSIP1 | 0.0604733 | -0.504228 |
| seq.23366.15 | Ubiquitin-like protein 7 | UBL7 | 0.0604733 | -0.360707 |
| seq.31106.18 | Cdc42 effector protein 3 | CDC42EP3 | 0.0604733 | -1.989699 |
| seq.12709.63 | Histone H1x | H1-10 | 0.0604733 | -1.383834 |
| seq.5687.5 | Glucosidase 2 subunit beta | PRKCSH | 0.0604733 | -0.15956 |
| seq.9004.24 | Low-density lipoprotein receptor class A domain-containing protein 4 | LDLRAD4 | 0.0604733 | -0.643814 |
| seq.23601.43 | Acyl-coenzyme A thioesterase 12 | ACOT12 | 0.0604733 | -0.377842 |
| seq.32648.65 | ATP-dependent RNA helicase DDX55 | DDX55 | 0.0604733 | -0.578023 |
| seq.31362.50 | DNA-binding protein A | YBX3 | 0.0604733 | -1.296323 |
| seq.21891.31 | Fibulin-7 | FBLN7 | 0.0604733 | 0.2501456 |
| seq.9169.14 | Small ubiquitin-related modifier 3 | SUMO3 | 0.0604733 | -0.320931 |

| | | | | |
|---|---|---|---|---|
| seq.31447.34 | High mobility group protein B1 | HMGB1 | 0.0604733 | -0.8725 |
| seq.15331.47 | Histone-binding protein RBBP4 | RBBP4 | 0.0604733 | -1.067952 |
| seq.6294.11 | B melanoma antigen 2 | BAGE2 | 0.0604733 | -0.65495 |
